# Travelling wave and asymptotic analysis of a multiphase moving boundary model for engineered tissue growth

**DOI:** 10.1101/2021.10.06.463324

**Authors:** Jacob M. Jepson, Nabil T. Fadai, Reuben D. O’Dea

## Abstract

We derive a multiphase, moving boundary model to represent the development of tissue *in vitro* in a porous tissue engineering scaffold. We consider a cell, extra-cellular liquid and a rigid scaffold phase, and adopt Darcy’s law to relate the velocity of the cell and liquid phases to their respective pressures. Cell-cell and cell-scaffold interactions which can drive cellular motion are accounted for by utilising relevant constitutive assumptions for the pressure in the cell phase. We reduce the model to a nonlinear reaction-diffusion equation for the cell phase, coupled to a moving boundary condition for the tissue edge, the diffusivity being dependent on the cell and scaffold volume fractions, cell and liquid viscosities, and parameters that relate to cellular motion. Numerical simulations reveal that the reduced model admits three regimes for the evolution of the tissue edge at large-time: linear, logarithmic and stationary. Employing travelling wave and asymptotic analysis, we characterise these regimes in terms of parameters related to cellular production and motion. The results of our investigation allow us to suggest optimal values for the governing parameters, so as to stimulate tissue growth in an engineering scaffold.

## 1 Introduction

*In vitro* tissue engineering is a form of regenerative medicine which often involves seeding cells into a porous bio-engineered scaffold that allows nutrient transport, structural support and a means for cell signalling activity (Chan and Leong, 2008). Subject to the correct environment and growth factors, the cells will develop into a functional construct that can be used to restore damaged tissues and improve concepts in pharmaceutical research such as experimental drug therapy (Jensen et al., 2018). Employing contributions from an assortment of scientific fields, tissue engineering is considered an interdisciplinary practice that has the potential to benefit a substantial proportion of the global population with devastating soft tissue, bone and whole organ diseases (Dzobo et al., 2018). The field of tissue engineering has enjoyed many successes; for example, the generation, replacement and longevity of engineered bones and bronchial tubes derived from the recipients’ cells (Sato et al., 2008; Petite et al., 2000; Schimming and Schmelzeisen, 2010). However, a shortage in the supply of donor tissue creates a demand on the field to make engineered tissue routinely clinically available (Levitt, 2015). Whilst the field is rich in both theoretical and experimental knowledge, a lack of understanding regarding the processes by which cells assemble into tissues means that viable replacement constructs are only available in a minority of cases.

Extensive mathematical research has been undertaken to make sense of the complicated mechanisms within tissue growth. Some authors adopt a microscale approach, which can take the form of cellular automaton systems (Vitvitsky, 2014; Lehotzky and Zupanc, 2019; Youssef, 2015) that seek to model interactions between a large number of individual cells. Whilst such systems can track the behaviour of cells, they can become computationally infeasible for tissue-scale simulations (Ermentrout and Edelstein-Keshet, 1993). Some authors adopt a probabilistic approach (Fadai et al., 2019; Browning et al., 2019a,b) and exploit experimental data to explore ways in which tissue engineering techniques can be improved. For example, Sogutlu and Koc (2013) present a stochastic model to determine the expected required number of pores for each region of a tissue engineering scaffold.

Conversely, continuum models (O’Dea et al., 2010; Lemon et al., 2006; Breward et al., 2002; Eyles et al., 2019; Byrne et al., 2003) track the evolution of tissue constituents by employing systems of partial differential equations. Whilst continuum models cannot impose rules on individual cells, they can be derived by imposing averaging techniques (Drew, 1983) on equations that govern cellular behaviour at a microscopic level, from which relevant mathematical techniques may be exploited to determine relationships between parameters and model behaviours. The study of *in vitro* tissue growth via continuum models is extensive, see O’Dea et al. (2012) and Klika et al. (2016) for reviews; of particular relevance to this study however, Lemon et al. (2006) considers a continuum multiphase model to investigate how mechanical pressures within growing tissue influence the aggregation or dispersion of cells in a scaffold, which relates the existence of these regimes to the governing parameters. Lemon and King (2007a) examine travelling wave solutions of the multiphase model formulated in Lemon et al. (2006), and find that in certain limits, the tissue propagates through the scaffold at a constant speed as either a forward or backward travelling wave, dependent on parameter values.

In this paper, we develop and analyse a continuum multiphase model that represents the development of tissue *in vitro* in an artificial scaffold. In our model, we aim to capture key features of tissue growth whilst developing a tractable formulation; in particular, we consider a porous flow description comprising a tissue cell phase, extra-cellular liquid phase and a scaffold phase, the former two being modelled as incompressible fluids and the latter as an inert solid. Tissue mechanics are accounted for by considering relevant constitutive assumptions in a similar fashion to those presented in Lemon et al. (2006) and Lemon and King (2007a). The model is reduced to a reaction-diffusion equation for the cell phase and a moving boundary condition for the tissue edge, after which travelling wave, asymptotic and numerical methods are employed to deduce the resulting solution behaviour. The paper is constructed as follows. In Section 2, we formulate and subsequently reduce and non-dimensionalise the model. In Section 3, we present and discuss numerical solutions to the reduced model, which motivate the travelling wave and asymptotic analyses conducted in Sections 4 and 5. In Section 6, we draw some conclusions regarding the behaviour of the model and interpret the mathematical results in terms of the biological application.

## 2 Model development

We construct a multiphase model to describe the growth of a nutrient rich tissue within a porous tissue engineering scaffold. For simplicity, we formulate the model in a one-dimensional Cartesian geometry. The model consists of three phases: two of which are fluid phases denoted by *n*(*x, t*) and *w*(*x, t*), and represent the volume fraction of cells and extra-cellular liquid, respectively. A rigid, non-degradable scaffold with uniform volume fraction *s* is the third phase and remains constant and uniform, the porosity of the scaffold hence being given by 1 − *s*. Cell growth and death occur via mass transfer between *n* and *w*. The phases satisfy the no voids volume constraint:

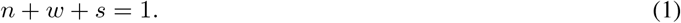

The velocity fields *v*_*n*_(*x, t*) and *v*_*w*_(*x, t*), and pressures *p*_*n*_(*x, t*) and *p*_*w*_(*x, t*), are associated with the phases *n* and *w* accordingly. The spatial domain of the tissue evolves over time due to cellular motion, so we track it with a moving boundary, *x* = *L*(*t*). In the subsections that follow, we state equations that govern mass transfer between *n* and *w*, as well as provide constitutive assumptions for *v*_*n*_, *v*_*w*_, *p*_*n*_ and *p*_*w*_ suitable to describe tissue growth in a scaffold. We state necessary initial and boundary conditions for the variables and the moving boundary *L*(*t*), and simplify and non-dimensionalise the model.

### 2.1 Governing equations

We assume that cells proliferate and assemble daughter cells from the available liquid, and that when cells die, they decompose and dissolve into the liquid phase. In view of these processes, it is reasonable to follow Lemon et al. (2006), Byrne et al. (2003), Breward et al. (2002) and Preziosi and Tosin (2003) (and many others) and assume the densities of *n* and *w* to be equal. Following these assumptions, the mass transfer equations can be represented as

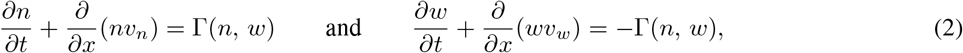

where Γ is the net rate of cell proliferation. Adding the equations from (2) results in the overall conservation of mass condition

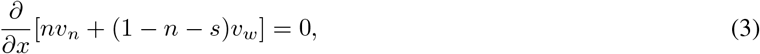

where (1) has been used to eliminate the time derivative and to replace *w* with 1 − *n* − *s*.

Noting that *n* and *w* are modelled as fluids and *s* as a porous scaffold, we take the interphase drags to be dominated by those with the scaffold and neglect that between the tissue and liquid. In view of this, we apply Darcy’s law to relate the velocity of the cell and liquid phases to their respective pressures. Following King and Franks (2004) and Eyles et al. (2019), we take

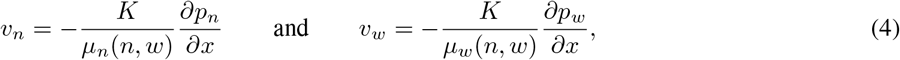

where *µ*_*n*_ and *µ*_*w*_ represent the viscosity of the cell and liquid phases and *K* is the permeability of the scaffold.

Remaining consistent with Lemon et al. (2006) and Lemon and King (2007a,b), we relate the cellular and extra-cellular liquid pressures via

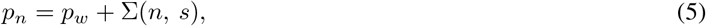

where Σ represents extra pressures that arise due to cell-cell and cell-scaffold interactions. Since the scaffold is assumed to be inert and of uniform porosity, we suppress the dependence Σ has on *s* from hereon for brevity. We note that combining (5) with the relations from (4) allows the elimination of *p*_*n*_ and *p*_*w*_ and provides

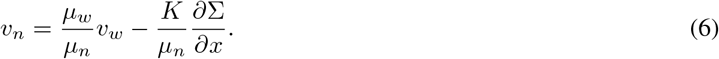

### 2.2 Initial and boundary conditions

Assuming the tissue to be symmetric about its centre (*x* = 0), we take

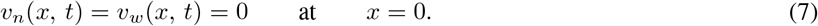

Naturally, the cell volume fraction is identically zero at the edge of the tissue:

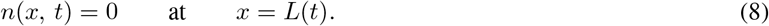

The moving boundary *L*(*t*) moves with the cell velocity, hence

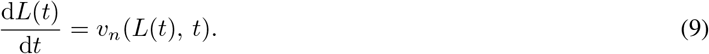

The initial distribution of *n* and tissue boundary position respectively are denoted by

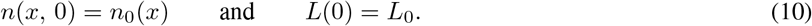

### 2.3 Model reduction

We reduce the model to a reaction-diffusion equation and a moving boundary condition. Integrating (3) and applying the boundary conditions from (7) provides

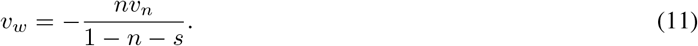

Combining (11) with (6) eliminates *v*_*w*_ and provides

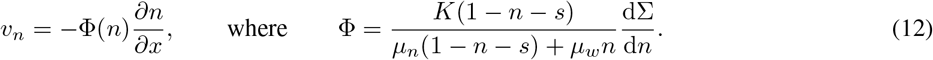

Here, we note *µ*_*n*_ and *µ*_*w*_ are assumed to be independent of *n* for simplicity. Substituting (12) into the first of (2) provides the reaction-diffusion equation:

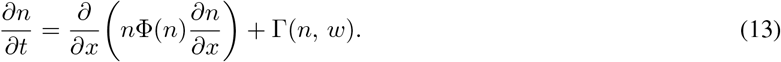

Combining (9) with (12) provides the moving boundary condition:

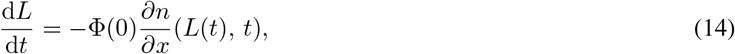

where the boundary condition from (8) provides Φ(*n*) = Φ(0) at *x* = *L*(*t*). Finally, (12) implies the boundary condition on *v*_*n*_ from (7) becomes

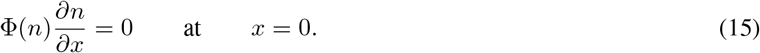

### 2.4 Constitutive assumptions

We now define constitutive assumptions for Γ and Σ that are suitable to describe tissue growth in a rigid scaffold. We assume that daughter cells are constructed via mitosis using the available liquid, and that when cells die via apoptosis they dissolve into the liquid. Thus, we take

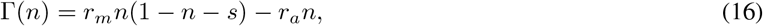

where *r*_*m*_ and *r*_*a*_ are the positive constant rates of cell mitosis and apoptosis, and (1) is used to replace *w* with 1 − *n* − *s*. Following Lemon et al. (2006) and Lemon and King (2007a,b), an appropriate expression for Σ(*n*) is

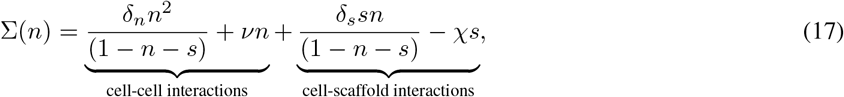

for *ν* ∈ ℝ and positive constants *δ*_*n*_, *δ*_*s*_ and *χ*. The first term in (17) represents repulsive forces exerted between the cells at high volume fractions, as characterised by the singularity at *n* = 1− *s*. The second term represents the propensity for cells to disperse or aggregate, with *ν* taking a positive or negative value accordingly. The third term represents repulsive forces that occur due to cell-scaffold interactions, whilst the fourth describes attractive forces between the cells and scaffold. For simplicity, we take *δ*:= *δ*_*n*_ = *δ*_*s*_. We note that Φ(*n*) must be strictly positive to prevent negative diffusion in (13) and nonlinear degeneracy in (14). This is achieved when *ν* > 0, which is henceforth assumed. Physically, this corresponds to a tendency for cells to spread through the scaffold (Lemon et al., 2006).

### 2.5 Non-dimensionalisation and parameter values

We non-dimensionalise (13), (14) and the initial and boundary conditions from (8), (10) and (15). By introducing the dimensionless variables

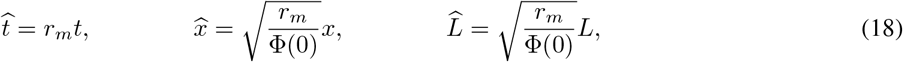

the following dimensionless model results:

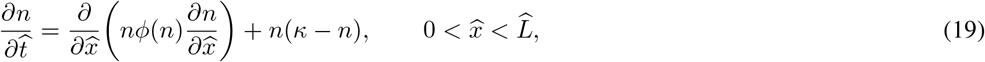

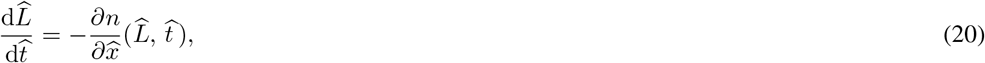

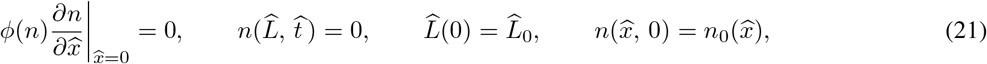

where *κ* = 1 − *s* − *r*_*a*_*/r*_*m*_ and 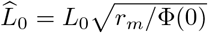. We also have

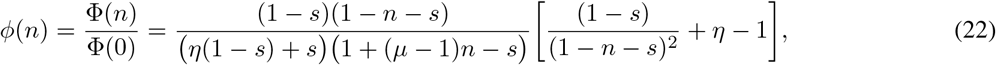

where *µ* = *µ*_*w*_*/µ*_*n*_ and *η* = *ν/δ*. In the proceeding, we dispense of the hat notation for clarity.

A linear stability analysis around the steady states of (19), *n* = 0 and *n* = *κ*, is conducted in Appendix A and shows that the former is stable when *κ* < 0 and the latter when *κ* > 0. In view of this, we primarily investigate (19)–(21) for different values of *κ*, though variations in *s, µ*, and *η* will also be considered in part so as to deduce their optimal values for the stimulation of tissue growth. Unless otherwise stated, we take *µ* = *η* = 1 and adopt the initial conditions

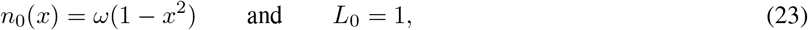

so that *ω* denotes the cell volume fraction at *x* = 0. Following Lemon and King (2007a,b), and unless otherwise stated, we set *s* = 0.2 and *ω* = 0.03, the former corresponding to a scaffold with a porosity of 0.8 and is consistent with the experimental study presented in Malda et al. (2004).

## 3 Numerical results

We present and discuss the numerical solutions for *n*(*x, t*) and *L*(*t*) from the PDE system (19)–(21), paying separate attention to the cases *κ* > 0, *κ* < 0 and *κ* = 0. For numerical convenience, we fix the moving boundary by introducing the variable transform *ξ* = *x/L*(*t*) so that *ξ* ∈ [0, 1], which means (19) and (20) become

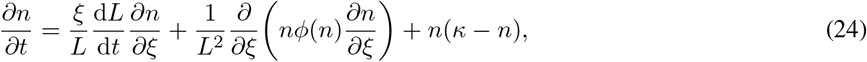

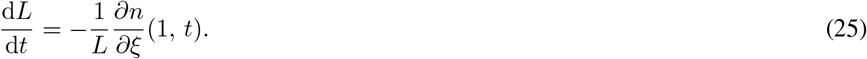

Subject to the transformed boundary and initial conditions from (21), we numerically integrate (24) and (25) by discretising first and second order spatial derivatives using second order finite differences. Upwind finite differences were used for the second term of (24). Temporal derivatives are numerically integrated by utilising `ode23s` in MATLAB.

For *κ* = 0.3, as seen in Figure 1(a, b), we observe semi-infinite travelling waves in *n* and linear growth in *L* after a period of transient growth from their initial states. Numerical simulations that are not included here suggest that travelling wave behaviour is exhibited by (19)–(21) for all *κ* > 0. A travelling wave and asymptotic analysis of (19)–(21) for *κ* > 0 and 0 < *κ* ≪ 1 are discussed in Sections 4 and 5 accordingly.

**Figure 1:**
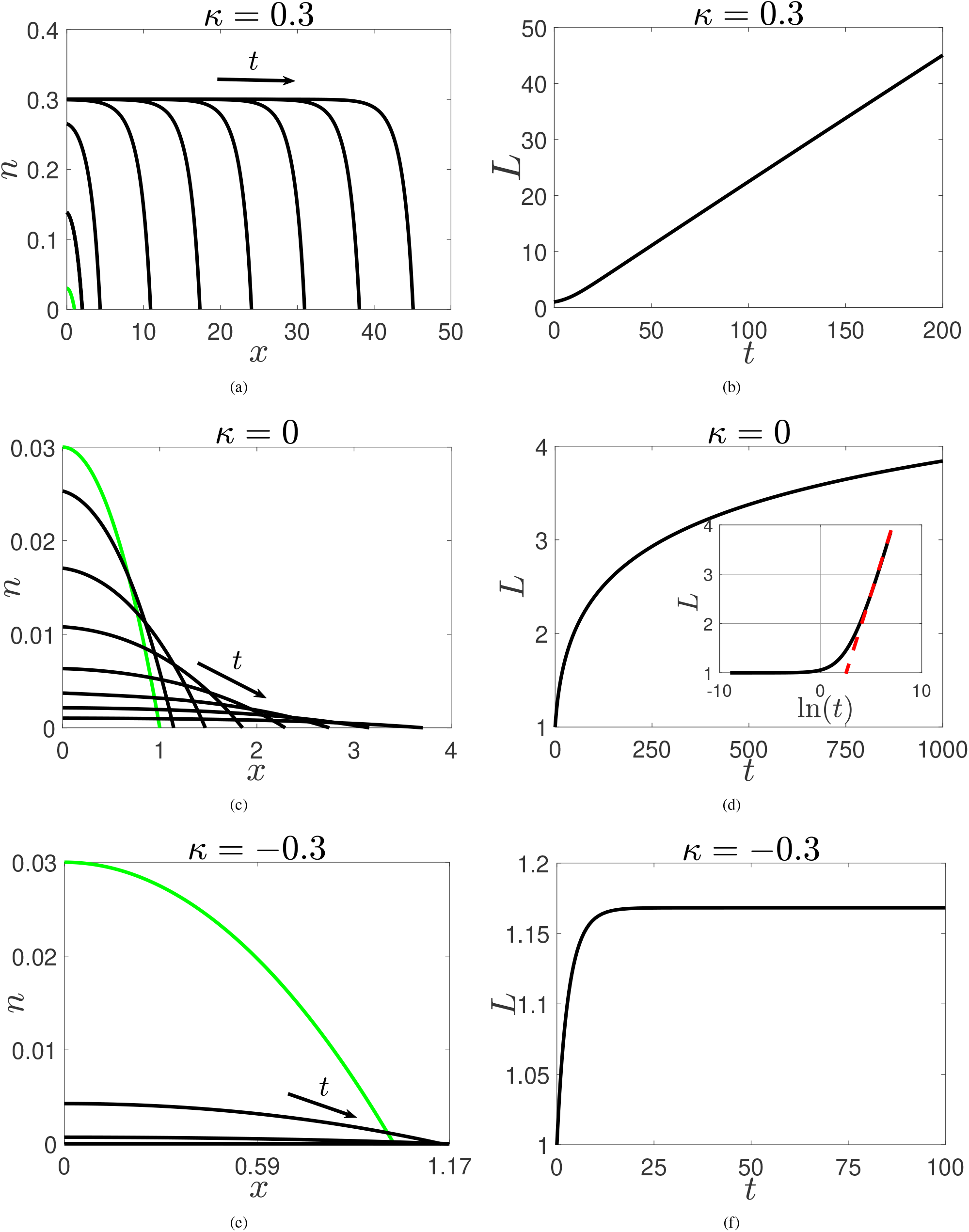
Numerical solutions of the system of PDEs from (19)–(21). The green lines in (a, c, e) represent the initial conditions *n*_0_(*x*) from (23), whereas the black lines represent *n* for different values of *t*. The black arrows point in the direction of increasing time. The dashed red line in (d) highlights the logarithmic growth of *L* at large-time. Parameter values: *L*_0_ = *µ* = *η* = 1, *ω* = 0.03 and *s* = 0.2

For *κ* = 0, as seen in Figure 1(c), we observe *n* decaying from the initial data. Figure 1(d) shows unbounded growth in *L*. The inset shows *L* plotted against ln(*t*), from which we conclude that *L* grows logarithmically at large-time. An asymptotic analysis of (19)–(21) for *κ* = 0 is undertaken in Section 5.

For *κ* = −0.3, as seen in Figure 1(e), we observe that *n* decays from the initial data more quickly than for *κ* = 0. The initial growth of *L* shown in Figure 1(f) occurs due to the diffusion of *n* from the initial state; however, we observe the eventual formation of a steady state. In accord with expectations, we note that *n* is non-zero at large-time when *κ* < 0 since *n* = 0 is a steady state of (19). Numerical simulations that are not included here suggest the steady state behaviour in *L* and decay in *n* is exhibited by (19)–(21) for all *κ* < 0. An asymptotic analysis of (19)–(21) is carried out for *κ* < 0 when |*κ*| ≪ 1 in Section 5.

## 4 Travelling wave analysis for *κ* > 0

Figures 1(a,b) indicate the emergence of semi-infinite travelling waves of constant speed for *κ* > 0. In light of this, we assume that for sufficiently large-time, *L* ∼ *ct* where *c* is the constant wave speed. In this section, we employ travelling wave analysis to obtain the wave speed *c* in terms of the governing parameters for *κ* > 0.

We transform (19)–(21) via the travelling wave coordinates *z* = *x* − *L* ∼ *x* − *ct* where *z* ∈ (−∞, 0]. Setting *n*(*x* − *ct*) = *n*(*z*), we obtain

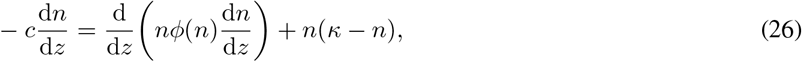

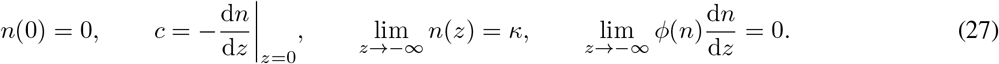

Following Fadai and Simpson (2020) and Fadai (2021), we define

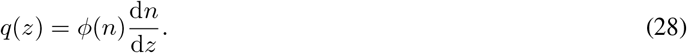

Multiplying (26) by *ϕ*(*n*) and re-writing the conditions from (27) in terms of *q*(*z*), we obtain

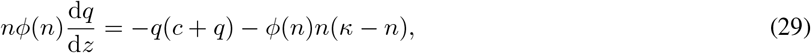

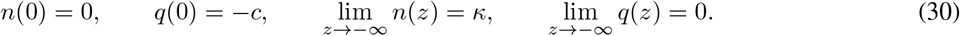

Here, we note the second boundary condition from (27) transforms into the second of (30) because *ϕ*(0) = 1. Dividing by (28) we have

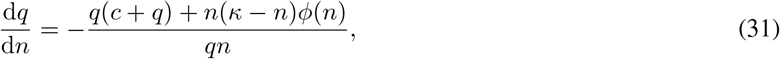

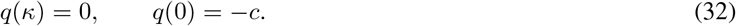

Using the shooting method (presented in Appendix B) to find the heteroclinic connection *q*(*n*) that connects (*n, q*) = (*κ*, 0) to (0, −*c*), we can determine a numerical approximation of the wave speed in terms of the governing parameters *κ, s, µ* and *η*.

In Figure 2(a,b), the solid black line represents the relationships *c*(*κ*) and *c*(*s*) respectively when (31)–(32) is approximated by the shooting method. The dashed green line represents these wave speeds when obtained by numerically solving (19)–(21), and computing *c* by evaluating d*L/*d*t* at large-time. In view of the close agreement between these two approaches to computing *c*, we henceforth concentrate on solutions provided by (31)–(32) for simplicity.

**Figure 2:**
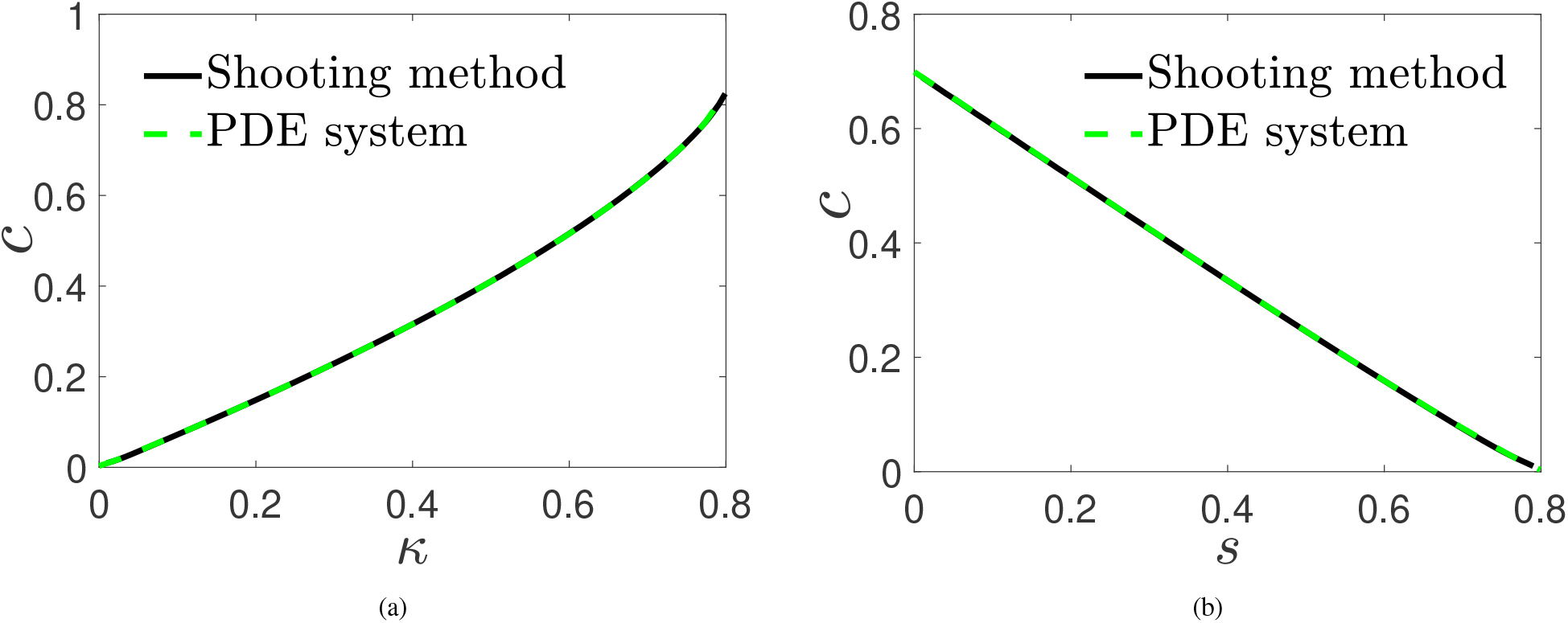
Numerical solutions for the wave speeds *c*(*κ*) and *c*(*s*). The solid black and dashed green lines represent approximations sought by numerically solving (31)–(32) using a shooting method and the system from (19)–(21), respectively. Parameter values for (a): *µ* = *η* = *L*_0_ = 1, *s* = 0.2 and *ω* = 0.03. Parameter values for (b): *µ* = *η* = *L*_0_ = 1, *r*_*a*_*/r*_*m*_ = 0.2 and *ω* = 0.03

The results presented in Figure 2 suggest that larger *κ* and smaller *s* increase the speed at which the tissue front grows, and since *κ* is dependent on *s*, this further corresponds to minimising *r*_*a*_*/r*_*m*_ and maximising the porosity of the scaffold. In Figure 3(a, b) we present the wave speeds *c*(*µ, η*) for *κ* = 0.3 and *κ* = 0.7, respectively. These results suggest that, for a fixed value of *κ*, the wave speed is maximised when *µ, η* → 0. Physically, this corresponds to the case where the viscosity of the cells dominates that of the liquid, and where repulsive forces exerted due to cell-cell and cell-scaffold interactions dominate inter-cellular forces that give rise to cell dispersal. Furthermore, Figure 3(a, b) indicates that the dependence *c* has on *µ* is weaker for *κ* = 0.3 than *κ* = 0.7. This suggests that for smaller *κ*, cell-cell and cell-scaffold interactions which can drive cellular motion are more prominent in controlling the wave speed than the cell and liquid viscosities. Additionally, and in agreement with Figure 2(a), Figure 3 indicates that the wave speed increases as *κ* increases.

**Figure 3:**
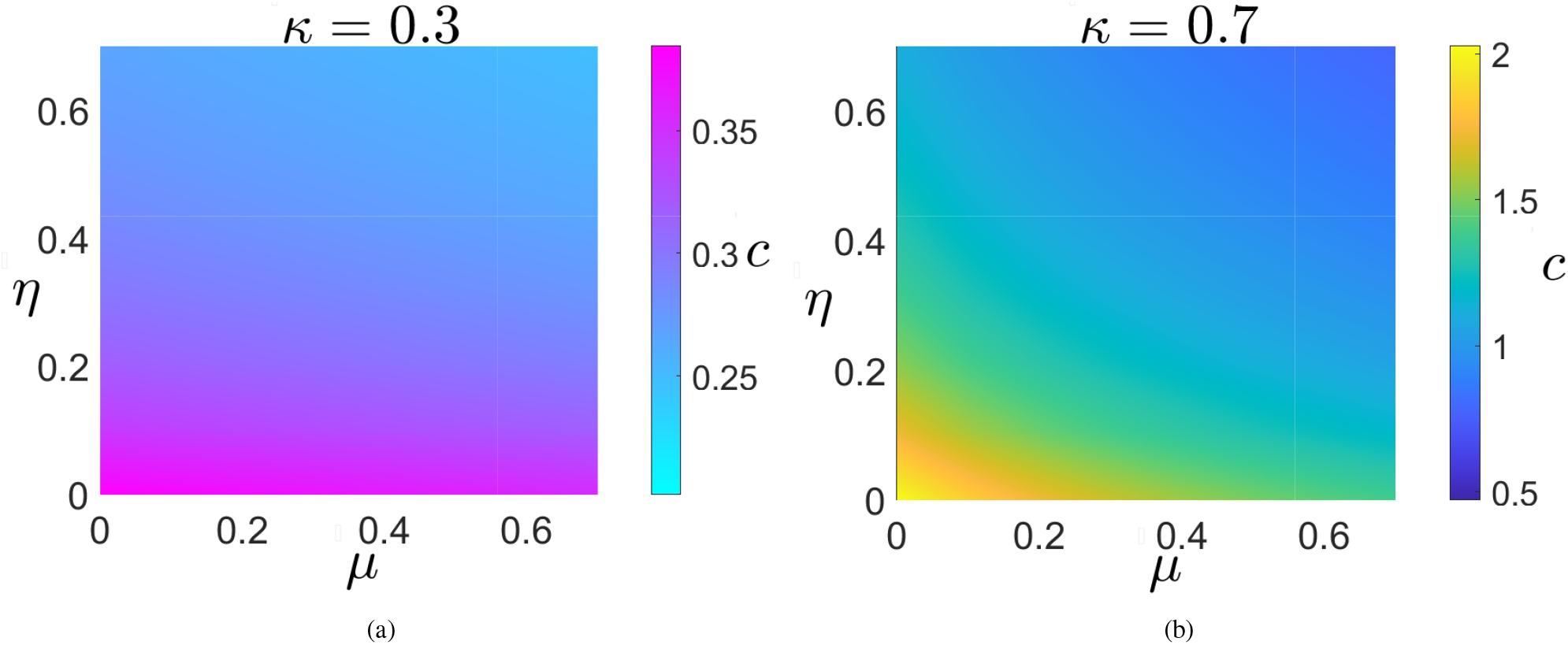
Numerical solutions for the wave speeds *c*(*µ, η*) for *κ* = 0.3 and *κ* = 0.7 sought by numerically solving (31)–(32) using a shooting method. *s* = 0.2 was used for both sub-figures. We note that the colour axes are different in each sub-figure.

## 5 Asymptotic analysis for *κ* = 0 and |*κ*| ≪ 1

In this section, we construct asymptotic solutions for *n*(*x, t*) and *L*(*t*) for *κ* = 0 and |*κ*| ≪ 1. The numerical results in Figures 1(c, e) indicate that after transient growth from the initial data, *n* ≪ 1 holds for both *κ* < 0 and *κ* = 0. Furthermore, given that *n* = *κ* is stable for *κ* > 0, *n* ≪ 1 is also expected when 0 < *κ* ≪ 1 for sufficiently large-time, hence *ϕ*(*n*) ∼ *ϕ*(0) = 1 is exploited in this section for all of these cases. To aid the subsequent asymptotic analysis, we introduce the variables *n* = *e*^*κt*^*N* (*x, T*) and *T* = (*e*^*κt*^ − 1)*/κ*, which imply (19) is simplified to

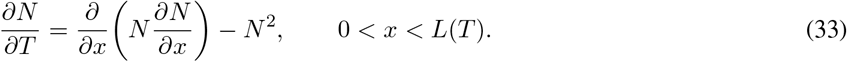

To analyse the behaviour of this equation, we follow Newman (1980) and adopt the ansatz

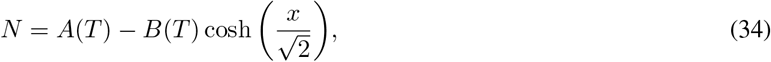

wherein 0 < *B* < *A* ≪ 1. Imposing *N* (*L, T*) = 0 on (34), we obtain

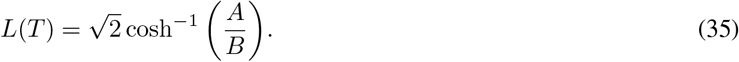

In view of (34), (33) provides

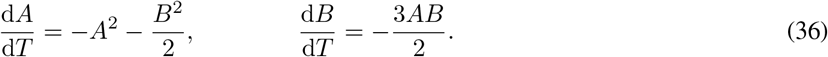

In this section, initial conditions for *N* and *L* are chosen to satisfy (34) and (35) when *T* = 0 – i.e., *A*(0) = *A*_0_, *B*(0) = *B*_0_ and hence 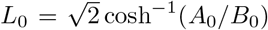. An implicit solution for *N* and *L* is found by computing *A*(*B*) and exploiting the relation from (35). Following Newman (1980), we have

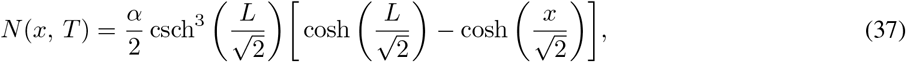

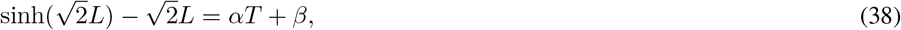

where 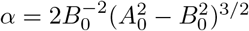and 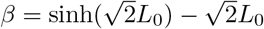.

We now deduce the large *T* behaviour of *N* (*x, T*) and *L*(*T*), from which the large-time behaviour of *n*(*x, t*) and *L*(*t*) when *κ* = 0 and |*κ*| ≪ 1 can subsequently be determined. Guided by the numerical results from Figure 1(d), the evolution of *L* corresponding to (33) satisfies *L* ≫ *L*_0_ for sufficiently large *T*, so we have from (38) that

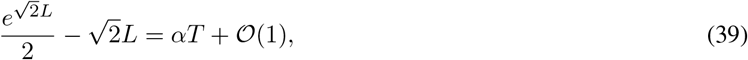

which is then inverted to give

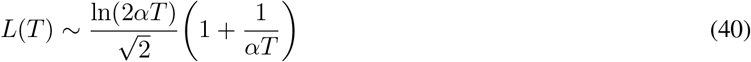

for *T* ≫ 1. Equation (37) and the leading-order term in the above expansion are used to find the following large *T* approximation for *N* :

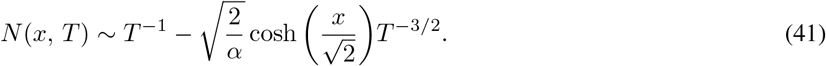

The large-time behaviour for *N* and *L* characterised by (40) and (41) are only valid for initial conditions that satisfy (34) and (35). The broader implications for (33) for a wider class of initial data, such as those employed in (23), are as follows. The asymptotic behaviour of (33) as *T* → ∞ comprises an interior layer near the interface within which the similarity reduction *N* ∼ *T* ^−1^*f* (*θ*) holds where *θ* = *x* − *L* and *L* = *b* ln(*T*), so that *f* (*θ*) satisfies

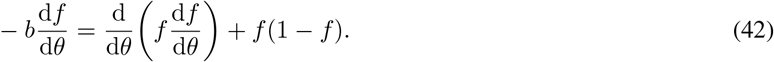

Following familiar arguments to that of the Porous Fisher Equation (Aaronson, 1980; Murray, 2002), although we emphasise that *N* is not a travelling wave of the usual form, the solution to (42) is given by

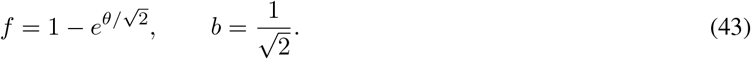

For *x* = 𝒪(1), setting *N* = *T* ^−1^ + *G*(*x, T*) implies

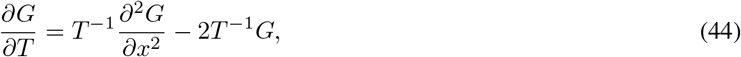

the solution to which that matches into the exponential terms in (43), and hence the corresponding term in the interior layer is given by

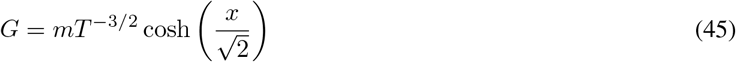

for some constant *m*, and this dominates the asymptotic behaviour of (44) as *T* → ∞. We now exploit (41) and (40) to deduce the large-time behaviour of *n*(*x, t*) and *L*(*t*) when *κ* = 0 and |*κ*| ≪ 1.

### 5.1 *κ* = 0

The large-time behaviour for *n*(*x, t*) and *L*(*t*) when *κ* = 0 can be extracted directly from (40) and (41) given that *N* = *n* when *κ* = 0 and lim_*κ*→0_ *T* = *t*. In Figure 4, we compare the numerical solution for *n* and *L* when obtained by numerically solving the PDE system from (19)–(21) for *κ* = 0 against their respective asymptotic solutions from (40) and (41). Overall, we observe an excellent agreement between the numerical and asymptotic solutions. We now characterise the large-time behaviour of *n*(*x, t*) and *L*(*t*) when |*κ*| ≪ 1.

**Figure 4:**
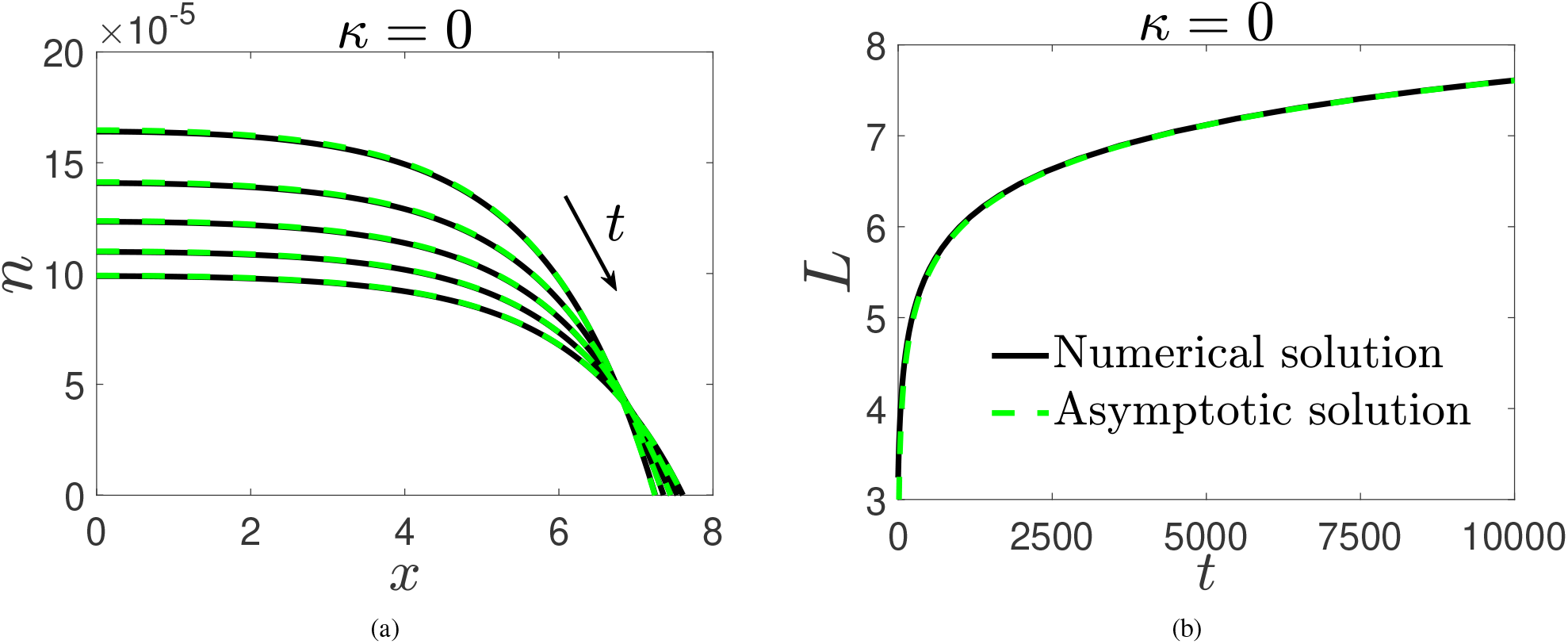
Numerical solution (solid black line) of the PDE system from (19)–(21) vs asymptotic solution (dashed green line) from (40) and (41) for *n* (a) and *L* (b) for *κ* = 0. Solutions for *n* are presented on *t* ∈ [6000, 10000] at intervals of 1000. Initial conditions for the numerical simulations were chosen to satisfy (34) and (35). Parameter values: *µ* = *η* = 1, *s* = 0.2, *A*_0_ = 0.05 and *B*_0_ = 0.01

### 5.2 |*κ*| ≪ 1

When 0 < *κ* ≪ 1, then *n*(*x, t*) takes the form of a travelling wave of constant speed, hence *L* ≫ *L*_0_ for *t* ≫ 1 and (40) becomes

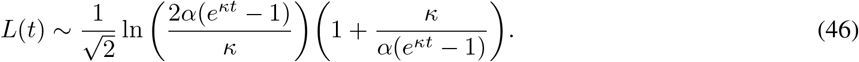

The leading-order logarithmic term in the above expansion and (37) are used to find the following large-time approxima-tion for *n:*

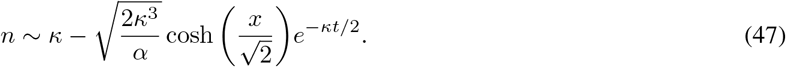

Equation (46) implies 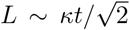 when *t* ≫ 1. Therefore, travelling waves propagate with speed 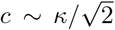 when 0 < *κ* ≪ 1, this being in agreement with the numerical results from Figure 2(a). We note that (46) and (47) hold for *κ* = 𝒪(1) if *µ* ≪ 1 and *η* ≫ 1 because *L* ≫ *L*_0_ and *ϕ*(*n*) ∼ 1 in this case. In Figure 5, we compare the numerical solution for *n* and *L* when obtained by numerically solving the PDE system from (19)–(21) for *κ* = 0.001 against their respective asymptotic solutions from (47) and (46). Overall, an excellent agreement between the numerical and asymptotic solutions is observed.

**Figure 5:**
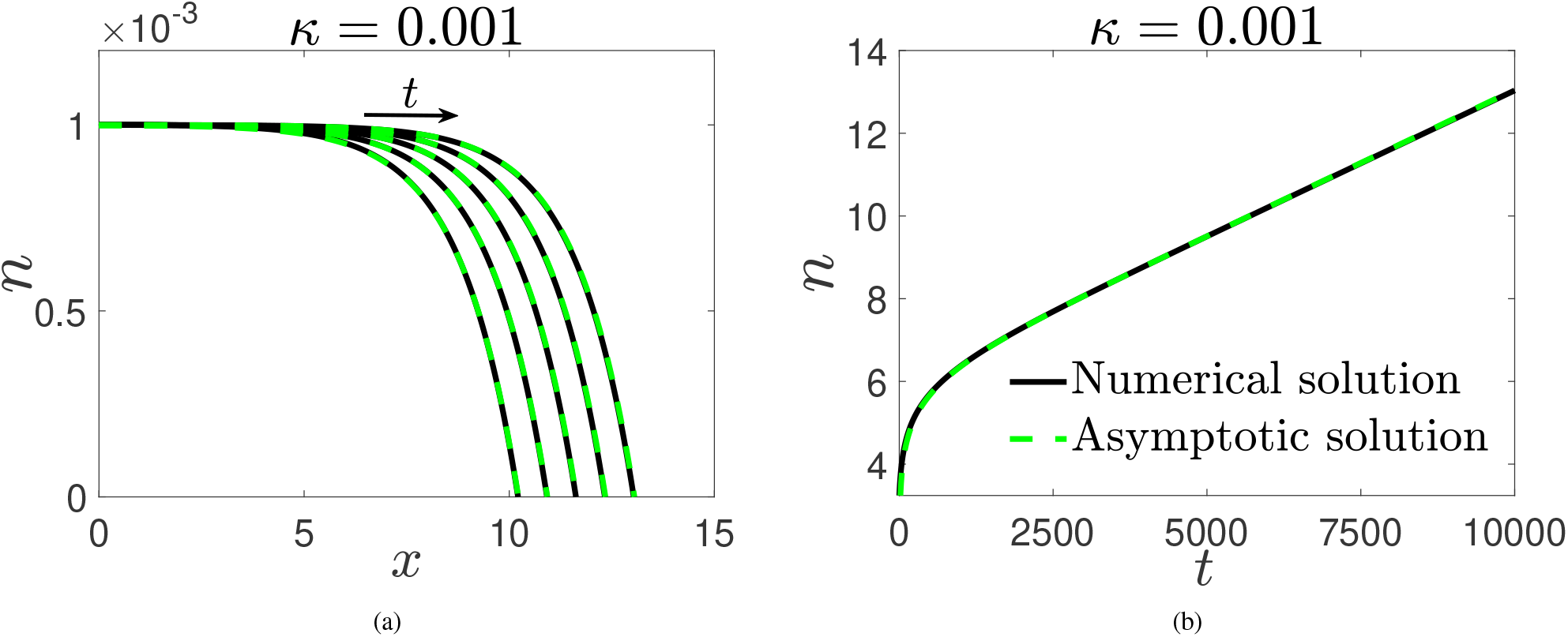
Numerical solution (solid black line) of the PDE system from (19)–(21) vs asymptotic solution (dashed green line) from (47) and (46) for *n* (a) and *L* (b) for *κ* = 0.001. Solutions for *n* are presented on *t* ∈ [6000, 10000] at intervals of 1000. Initial conditions for the numerical simulations were chosen to satisfy (34) and (35). Parameter values: *µ* = *η* = 1, *s* = 0.2, *A*_0_ = 0.05 and *B*_0_ = 0.01

When *κ* < 0, the numerical results in Figure 1(d) suggest that *L* → *L*_∞_ as *t* → ∞ for some finite constant *L*_∞_. In general, a large-time solution for *L* is unavailable given that *L*_∞_ ≫ *L*_0_ does not necessarily hold when *t* ≫ 1; however, when |*κ*| ≪ 1, then *κ* = 0 to leading order at *t* = 𝒪(1) and *L* evolves according to (40) until *t* = 𝒪(1*/*|*κ*|). In view of this, (46) is an appropriate asymptotic approximation for *L*(*t*) when *κ* < 0 and |*κ*| ≪ 1, the exponential terms being negligible at large-time and thus providing

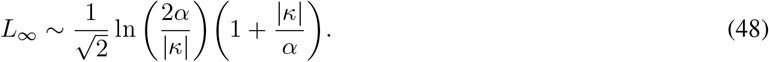

Combining (37) with the leading-order term from (46) provides the large-time approximation for *n* :

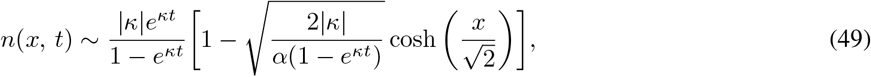

for *κ* < 0 and |*κ*| ≪ 1. In Figure 6, we compare the numerical solution for *n* and *L* when obtained by numerically solving the PDE system from (19)–(21) with *κ* = −0.001 against their respective asymptotic solutions from (49) and (46). Overall, a good agreement between the numerical and asymptotic solutions is observed. We note that for *κ* < 0 and |*κ*| = 𝒪(1), then (37) suggests that *n* = 𝒪(*e*^*κt*^) at large-time since *L*_∞_ = 𝒪(*L*_0_).

**Figure 6:**
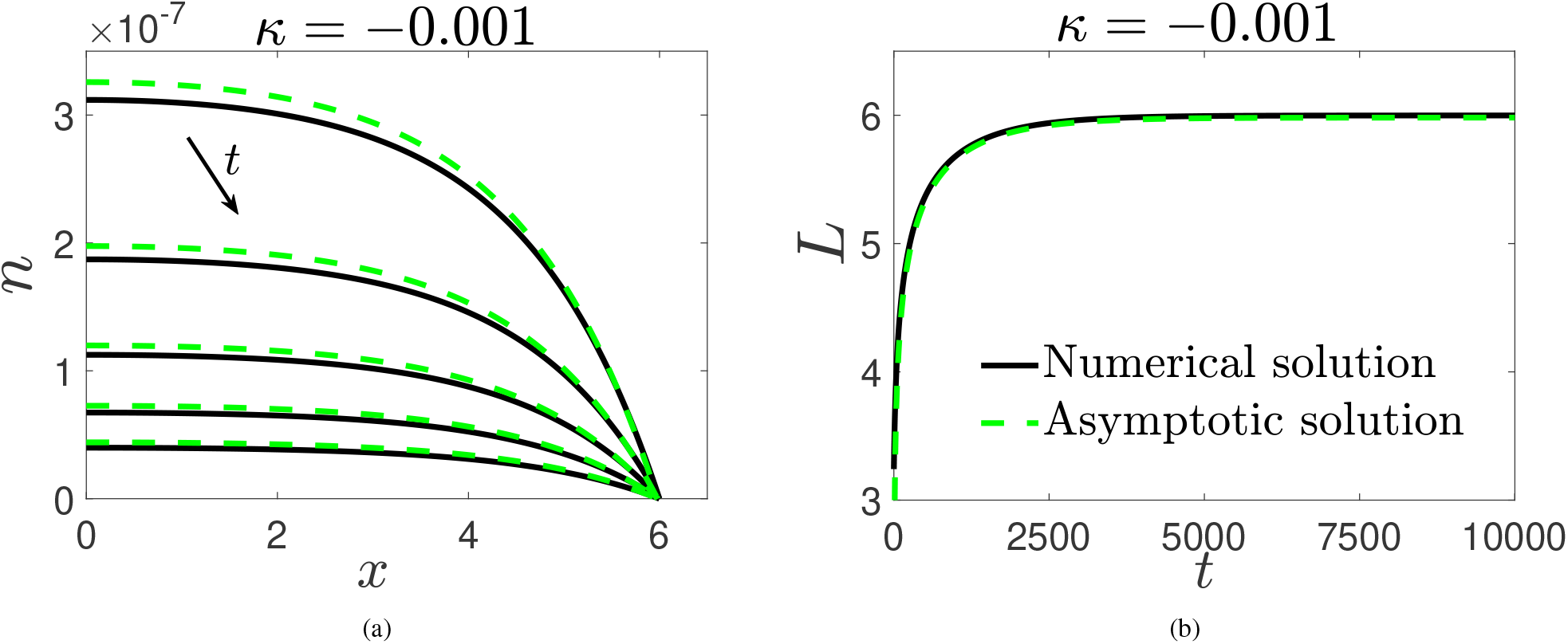
Numerical solution (solid black line) of the PDE system from (19)–(21) vs asymptotic solution (dashed green line) from (49) and (46) for *n* (a) and *L* (b) for *κ* = −0.001. Solutions for *n* are presented on *t* ∈ [8000, 10000] at intervals of 500. Initial conditions for the numerical simulations were chosen to satisfy (34) and (35). Parameter values: *µ* = *η* = 1, *s* = 0.2, *A*_0_ = 0.05 and *B*_0_ = 0.01

## 6 Conclusions

In this paper, we present a multiphase model to represent the development of tissue *in vitro* within a porous tissue engineering scaffold. We consider a tissue cell phase, extra-cellular liquid phase and a scaffold phase, and adopt Darcy’s law to relate the velocity of the cell and liquid phases to their respective pressures. The model includes mechanisms to represent cell growth and death, and pressures that arise from cell-cell and cell-scaffold interactions. We employ a moving boundary, *x* = *L*(*t*), to track the speed at which the tissue edge propagates through the scaffold. We reduce the model to a nonlinear reaction-diffusion equation for the cell volume fraction, *n*(*x, t*), and a moving boundary condition for the tissue edge. The diffusivity of the reaction-diffusion equation is dependent on the cell and scaffold volume fractions; cell and liquid viscosities, and pressures that arise from cell-cell and cell-scaffold interactions. One dimensionless parameter that is of crucial importance to the qualitative features of the cell phase evolution is *κ*, defined as difference between the scaffold porosity and the ratio between the cell death and growth rates. The model admits three regimes for the evolution of the cell volume fraction and the moving boundary, based on the sign of *κ*. Employing travelling wave and asymptotic analysis, we characterise these regimes in terms of *κ* and parameters related to cellular motion.

For *κ* > 0, we show that the cell volume fraction, *n*(*x, t*), spreads through the scaffold as a semi-infinite travelling wave with constant speed, emerging from the steady state *n* = *κ*. Employing travelling wave analysis, we accurately compute the wave speed (i.e., the speed at which the tissue edge propagates through the scaffold) as a function of the governing parameters. We find that the wave speed is greatest when the rate of apoptosis is negligible in comparison to that of mitosis; the scaffold volume fraction is small (and hence when the scaffold porosity is large); the viscosity of the cells dominate that of the liquid, and when repulsive forces exerted due to cell-cell and cell-scaffold interactions dominate inter-cellular forces that give rise to cell dispersal. Furthermore, we deduce that for smaller values of *κ*, cell-cell and cell-scaffold interactions which can drive cellular motion are more prominent in controlling the wave speed than the cell and liquid viscosities. Overall, the results obtained in this paper suggests that *κ* > 0 must hold for tissue growth to occur, thus suggesting that tissue engineers should ensure that the porosity of the scaffold is at least larger than the ratio between the rate of cell death and growth.

For *κ* = 0 and |*κ*| ≪ 1, we employ asymptotic analysis to find explicit solutions for *n* and *L*. For *κ* = 0, we conclude that the cell volume fraction no longer takes the form of a travelling wave, the leading order approximation for the cell volume fraction at the tissue centre being *n*(*x, t*) ∼ *t*^−1^ for *t* ≫ 1. We also show that the moving boundary grows logarithmically at large-time, a behaviour not frequently associated with tissue growth. For 0 < *κ* ≪ 1, travelling waves with speed 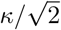 are observed in *n* for sufficiently large-time. The growth of the tissue edge is logarithmic until *t* = 𝒪(1*/κ*) and linear thereafter, thus suggesting the formation of travelling waves with constant speed is delayed as *κ* → 0^+^. For *κ* < 0, we deduce that the cell volume fraction decays exponentially with rate *κ* at large-time, with the moving boundary tending towards a steady state. For *κ* < 0 and |*κ*| ≪ 1, the evolution of the *L* is shown to be logarithmic until *t* = 𝒪(1*/κ*) and approaches a steady state thereafter, the value of which is found explicitly and related to *κ* and the initial conditions employed in the model. For *κ* = 0 and |*κ*| ≪ 1, an asymptotic analysis demonstrates that the initial aggregate of cells seeded within the scaffold does not influence the eventual large-time behaviour of the tissue or tissue edge.

For a functional tissue construct to develop within a scaffold, cells must be exposed to the correct environment and stimulated with growth factors such as oxygen. Whilst key features of tissue growth such as cell mitosis, apoptosis and motion are included in this paper, concepts such as environmental pressures and nutrient supply have not been considered. Therefore, following Lemon and King (2007a), a natural extension of this work would include examining the influence that nutrient limitation has on cell growth. We leave these extensions for future consideration.

## Data accessibility

The MATLAB codes employed to generate the results presented in all figures can be found at: https://github.com/JacobJepson/scaffoldcodes_2021

## Acknowledgments

Jacob M. Jepson acknowledges funding provided by the Engineering and Physical Sciences Research Council (grant number EP / R513283 / 1). We are grateful to Professor John King for inspiration for the work, and for significant support and guidance in its development.

## A Linear stability analysis

A linear stability analysis around the steady states of (19), *n*_∞_ = 0, *κ*, provides insight into the dependence of the model behaviour on *κ* and *ϕ*(*n*). Neglecting the influence of the moving boundary condition on the stability of (19), we linearise on a semi-infinite domain. We substitute *n* = *n*_∞_ + *ε* exp(*iγx* + *λt*) into (19) for a perturbation wave number *γ* and growth rate *λ* where *ε* ≪ 1. Considering terms of 𝒪(*ε*) only, the growth rate for perturbations of wave length 2*π/γ* is

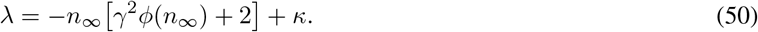

Since *ϕ*(*n*) is assumed to be positive for any *n*, the steady state *n*_∞_ = *κ* is stable for all *κ* > 0. For *n*_∞_ = 0, we have *λ* = *κ* which indicates stability when *κ* < 0.

## B The shooting method

In this section, we formulate a numerical shooting method to find the wave speed *c*, as stated in Section 4. To do this, we send trajectories of *q*(*n*) from (*κ*, 0) to find a trajectory that connects to (0, −*c*). Computationally, it is more straightforward to shoot trajectories forwards as opposed to backwards, and we hence introduce the change of variable *X* = *κ* − *n* and obtain

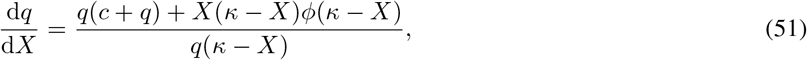

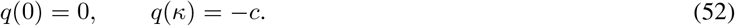

Noting that the end points are computationally singular, we use *q*(*ζ*) = −*ζ* and *q*(*κ* − *ζ*) = −*c* where 0 < *ζ* ≪ 1 is some user-defined tolerance. We employ the discrepancy function

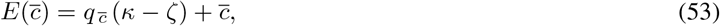

where 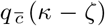 is the solution to (51) evaluated at *X* = *κ* − *ζ* for a trial wave speed 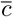. Equation (51) is numerically integrated with `ode23s` in MATLAB with the initial condition *q*(*ζ*) = −*ζ*. Using `fzero` in MATLAB to find the zero of 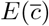, the wave speed *c* and the corresponding heteroclinic trajectory *q*(*X*) is determined.

## Notes

### Competing Interest Statement

The authors have declared no competing interest.

## References

D. G. Aaronson. Density-Dependent Interaction–Diffusion Systems. Dynamics and Modelling of Reactive Systems, Academic Press, 1980. 161–176.

C. J. W. Breward, H. M. Byrne, and C. E. Lewis. The role of cell-cell interactions in a two-phase model for avascular tumour growth. Journal of Mathematical Biology, 2002. 45:125–152.

A. P. Browning, P. Haridas, and M. J. Simpson. A Bayesian Sequential Learning Framework to Parameterise Continuum Models of Melanoma Invasion into Human Skin. Bulletin of Mathematical Biology, 2019a. 81:676–698.

A. P. Browning, S. W. McCue, R. N. Binny, M. J. Plank, E. T. Shah, and M. J. Simpson. Inferring parameters for a lattice-free model of cell migration and proliferation using experimental data. Journal of Theoretical Biology, 2019b. 437:251–260.

H. M. Byrne, J.R. King, D.L.S McElwain, and L. Preziosi. Two-Phase Model of Solid Tumour Growth. Applied Mathematics Letters, 2003. 16:567–573.

B. P. Chan and K. W. Leong. Scaffolding in tissue engineering: general approaches and tissue-specific considerations. European Spine Journal, 2008. 17(4): 467–479.

D. A. Drew. Mathematical modelling of two-phase flow. Annual Review of Fluid Mechanics, 1983. 15:261–291.

K. Dzobo, N. E. Thomford, D. A. Senthebane, H. Shipanga, A. Rowe, C. Dandara, M. Pillay, and K. S. C. M. Motaung. Advances in Regenerative Medicine and Tissue Engineering: Innovation and Transformation of Medicine. Stem Cells International, 2018. Article ID 2495848.

G. B. Ermentrout and L. Edelstein-Keshet. Cellular Automata Approaches to Biological Modeling. Journal of Theoretical Biology, 1993. 160:97–133.

J. Eyles, J. R. King, and V. Styles. A tractable mathematical model for tissue growth. Interfaces and Free Boundaries, 2019. 21(4): 463–493.

N. T. Fadai. Semi-infinite travelling waves arising in a general reaction–diffusion Stefan model. Nonlinearity, 2021. 34:725–743.

N. T. Fadai and M. J. Simpson. New travelling wave solutions of the Porous-Fisher model with a moving boundary. Journal of Physics A: Mathematical and Theoretical, 2020.53(9).

N. T. Fadai, R. E Baker, and M. J Simpson. Accurate and efficient discretizations for stochastic models providing near agent-based spatial resolution at low computational cost. Journal of the Royal Society Interface, 2019. http://doi.org/10.1098/rsif.2019.0421.

G. Jensen, C. Morrill, and Y. Huang. 3D tissue engineering, an emerging technique for pharmaceutical research. Acta Pharmaceutica Sinica B, 2018. 8(5): 756–766.

J. R. King and S. J. Franks. Mathematical analysis of some multi-dimensional tissue-growth models. European Journal of Applied Mathematics, 2004. 15: 273–295.

V. Klika, E. A. Gaffney, Y. Chek, and C. P. Brown. An overview of multiphase cartilage mechanical modelling and its role in understanding function and pathology. Journal of the Mechanical Behaviour of Biomedical Materials, 2016. 62:139–157.

D. Lehotzky and G. K. H. Zupanc. Cellular Automata Modeling of Stem-Cell-Driven Development of Tissue in the Nervous System. Developmental Neurobiology, 2019. 79(5):497–517.

G. Lemon and J. King. Travelling-wave behaviour in a multiphase model of a population of cells in an artificial scaffold. Journal of Mathematical Biology, 2007a. 24(1):57–83.

G. Lemon and J. R. King. Multiphase modelling of cell behaviour on artificial scaffolds: effects of nutrient depletion and spatially nonuniform porosity. Journal of Mathematical Medicine and Biology, 2007b. 24:5783.

G. Lemon, J. R. King, H. M. Byrne, O. E. Jensen, and K. M. Shakesheff. Mathematical modelling of engineered tissue growth using a multiphase porous flow mixture theory. Journal of Mathematical Biology, 2006. 52:571–594.

M. Levitt. Could the organ shortage ever be met? Life Sciences, Society and Policy, 2015. 11: 6.

J. Malda, T. B. F. Woodfield, F. V. D. Vloodt, F. K. Kooy, D. E Martens, J. Tramper, C. A V. Blitterswijk, and J. Rielse. The effect of PEGT/PBT scaffold architecture on oxygen gradients in tissue engineered cartilaginous constructs. Biomaterials, 2004. 25(26):5681–5796.

J. D. Murray. Mathematical Biology I: An Introduction. Springer-Verlag, 2002.

W. I. Newman. Some Exact Solutions to a Non-linear Diffusion Problem in Population Genetics and Combustion. Journal of Theoretical Biology, 1980. 85:325–334.

R. D. O’Dea, S. L. Waters, and H. M. Byrne. A multiphase model for tissue construct growth in a perfusion bioreactor. Mathematical Medicine and Biology, 2010. 27:95127.

R. D. O’Dea, H. M. Byrne, and S. L. Waters. Continuum Modelling of In Vitro Tissue Engineering: A Review. Computational Modeling in Tissue Engineering, 2012.

H. Petite, V. Viateau, W. Bensäid, A. Meunier, C. de Pollak, M. Bourguignon, K. Oudina, L. Sedel, and G. Guillemin. Tissue-engineered bone regeneration. Nature Biotechnology, 2000. 18(9):959–963.

L. Preziosi and A. Tosin. Multiphase modelling of tumour growth and extracellular matrix interaction: mathematical tools and applications. Journal of Mathematical Biology, 2003. 58:625–656.

T. Sato, H. Tao, M. Araki, H. Ueda, K. Omori, and T. Nakamura. Replacement of the Left Main Bronchus With a Tissue-Engineered Prosthesis in a Canine Model. The Annals of Thoracic Surgery, 2008. 86:422–428.

R. Schimming and R. Schmelzeisen. Tissue-engineered bone for maxillary sinus augmentation. Journal of Oral and Maxillofacial surgery, 2010. 62(6):724–729.

S. Sogutlu and B. Koc. Stochastic Modeling of Tissue Engineering Scaffolds with Varying Porosity Levels. Computer-Aided Design and Applications, 2013. 4:5 661–670.

A. A. Vitvitsky. Cellular automata with dynamic structure to simulate the growth of biological tissues. Numerical Analysis and Applications, 2014. 7:263–273.

B. B. Youssef. A parallel cellular automata algorithm for the deterministic simulation of 3-D multicellular tissue growth. Cluster Computing, 2015. 18:1561–1579.

